# Transcriptome Complexity Disentangled: A Regulatory Molecules Approach

**DOI:** 10.1101/2023.04.17.537241

**Authors:** Amir Asiaee, Zachary B. Abrams, Heather H. Pua, Kevin R. Coombes

## Abstract

Transcription factors (TFs) and microRNAs (miRNAs) are fundamental regulators of gene expression, cell state, and biological processes. This study investigated whether a small subset of TFs and miRNAs could accurately predict genome-wide gene expression. We analyzed 8895 samples across 31 cancer types from The Cancer Genome Atlas and identified 28 miRNA and 28 TF clusters using unsupervised learning. Medoids of these clusters could differentiate tissues of origin with 92.8% accuracy, demonstrating their biological relevance. We developed Tissue-Agnostic and Tissue-Aware models to predict 20,000 gene expressions using the 56 selected medoid miRNAs and TFs. The Tissue-Aware model attained an *R*^2^ of 0.70 by incorporating tissue-specific information. Despite measuring only 1/400th of the transcriptome, the prediction accuracy was comparable to that achieved by the 1000 landmark genes. This suggests the transcriptome has an intrinsically low-dimensional structure that can be captured by a few regulatory molecules. Our approach could enable cheaper transcriptome assays and analysis of low-quality samples. It also provides insights into genes that are heavily regulated by miRNAs/TFs versus alternative mechanisms. However, model transportability was impacted by dataset discrepancies, especially in miRNA distribution. Overall, this study demonstrates the potential of a biology-guided approach for robust transcriptome representation.

## 1. Introduction

The transcriptome comprises the entire set of messenger RNA molecules within an organism or cell and its regulation controls gene expression. Due to its ease of measurement, the transcriptome has played a significant role in advancing our understanding of differentiation [1–3], development [4–7], and disease [8–10]. The transcriptome is both an essential contributor to the cell’s underlying biology and a quantitative measure that provides a mathematical and concrete representation of the cell state [11,12].

Although RNA-Seq has become the most popular gene expression measurement technology due to the significant drop in its cost since 2008 [13], it is not always effective or feasible for measuring expression in biological or clinical specimens. For instance, when measuring gene expression on a scale of millions of experiments to determine cellular function or response (such as response to drugs or gene perturbations), the low cost per sample (ranging from USD 10 to USD 200, depending on the technology) might still be prohibitively expensive [14]. Additionally, when high-quality RNA samples are unavailable, alternative sensitive techniques like qPCR may be preferable, although they are not ideal for whole-transcriptome analysis due to the need for specific primers, limited throughput, and technical variability [15].

To address these challenges, the NIH LINCS project [14,16,17] developed a new, low-cost, high-throughput method for measuring a “reduced representation” of the gene expression profile consisting of 978 landmark genes and 80 constant control genes that together are called the L1000 assay. This method enables the inference of 11,350 non-measured transcripts with 81% accuracy, as measured by the correlation score. The authors employed this approach to scale the Connectivity Map project’s samples from 492 original samples [18] to more than 1.3 million samples [14], enabling the discovery of functional connections among diseases, genetic perturbations, and drug actions. In the case of low-quality RNA samples, the same set of predictive models can be theoretically used for predicting the transcriptome from the qPCR measurement of the L1000 genes.

Notably, the L1000 genes were chosen based on a pure, data-driven approach rather than prior biological knowledge, as highlighted by Subramanian et al. [14]. In short, their approach is based on the clustering of genes in the principal component space and selecting the centroids by an iterative peel-off procedure. Surprisingly, the authors reported that the landmark genes were not enriched in any specific functional class, such as transcription factors, as determined by hypergeometric overlap statistics [19].

The pure, data-driven selection method of predictive genes overlooks the causal relationships within the transcriptome, as some parts of the transcriptome act as the root causes or master regulators of others. For instance, certain transcripts produce transcription factor proteins that regulate the transcription of other genes [20]. Therefore the abundance of the former transcripts can be considered the cause of the latter ones. We hypothesize that taking into account the causal relationships between transcriptome components when making predictions can have two significant implications: firstly, by treating the causes as the main predictors for the entire transcriptome, the size of the “reduced representation” of the whole transcriptome could be substantially decreased; secondly, incorporating causal predictors into the model could result in more robust predictions across various environments. In this study, we use transcription factor (TFs) and microRNAs (miRNAs), two classes of regulatory molecules known to causally regulate transcription, as predictors to demonstrate that, by measuring only 28 of each class, we can explain 70% of the variation of the transcriptome, i.e., the expression of approximately 20,000 genes. Our findings suggest that the transcriptome has a low-dimensional structure that can be captured by measuring the expression of only 56 regulatory molecules. Each of our selected regulatory molecules represents a cluster of highly co-expressed regulatory molecules, which we term a regulatory component. Through gene enrichment analysis, we show that regulatory components consist of genes that are co-expressed and work together to control various biological processes [21,22], which is in contrast to the selected L1000 landmark genes. Additionally, we show that our approach requires two orders of magnitude fewer measurements than the traditional L1000 approach while achieving comparable whole-transcriptome prediction accuracy. This could lead to the development of cheaper transcriptome assays and accommodate low-quality RNA sample scenarios. Furthermore, we provide biological insights into the potential reasons for certain genes being easy or hard to predict using TFs and miRNAs by analyzing the sets of genes that are poorly or well explained by our model. Lastly, we evaluate the transportability of our model by assessing its performance in whole-transcriptome prediction based on three previously unseen datasets.

## 2. Results

### 2.1. Data Description

Before presenting our results, we first describe the datasets used in our analysis. Our analyses were performed on transcriptomic data from multiple sources, as described below.

#### 2.1.1. Main Dataset

The main data used in this study were collected from The Cancer Genome Atlas (TCGA) [23], a public data repository that contains clinical and omics data for over 10,000 patient samples from 32 different cancer types. To obtain the transcript data, we used the FireBrowse web portal and selected samples with matched mRNA and microRNA sequencing data. We obtained *n* = 8895 samples from 31 different cancer types. Glioblastoma, despite having a substantial number of samples, was excluded as almost all of its samples lacked the necessary microRNA measurements.

We downloaded the mRNA expression-level data that were quantified using the RSEM method [24] and normalized using FPKM-UQ (upper quartile normalized fragments per kilobase of transcript per million mapped reads). We obtained the microRNA expression data in the form of counts of mature microRNA normalized to reads per million mapped reads (RPM). We log2 transformed both the mRNA and miRNA datasets. We filtered the data by removing miRNAs with zero read counts in 75% of the samples, resulting in *p* = 470 remaining miRNAs [22]. Using the Transcription Factor Catalog [25], we identified 486 human transcription factors and separated them from the 20289 measured gene expressions [21]. The resulting dataset consisted of 8895 × (486 + 470) dimensions, comprising TFs and miRNAs as our original features and the rest of the transcriptome with 19803 gene expressions for the 8895 samples as outcomes.

The specific cancer types from TCGA that we used, along with their respective abbreviations, are presented in the Supplemental Data. It is important to understand that certain TCGA cancer types, such as Lung Adenocarcinoma and Lung Squamous-Cell Carcinoma, originate from the same tissue. They essentially represent different subtypes of a primary cancer based on that tissue’s lineage. When training our model on TCGA data and considering the cancer type labels, a more precise name for our model would be “Cancer Tissue Aware”. However, due to the broader applicability of our model beyond just TCGA and for the sake of simplified terminology, we use “Tissue Aware” as the descriptor for our model.

#### 2.1.2. Auxiliary Datasets

To ascertain the influence of normalization on our results, in addition to FPKM-UQ normalized data, we conducted all experiments using a TPM (Transcription Per Million)-normalized version of the TCGA data. Additionally, to evaluate the transportability of our whole-transcriptome prediction model trained on TCGA, we assessed its performance on three distinct datasets: GTEx (Genotype-Tissue Expression [26]), TARGET (Therapeutically Applicable Research to Generate Effective Treatments [27]), and the CCLE (Cancer Cell Line Encyclopedia [28]). These datasets were chosen due to their broad tissue and cell type coverage.

In detail, the CCLE offers an exhaustive genetic analysis of over 1000 human cancer cell lines, making it an invaluable repository for advancing new cancer treatments and probing the intricate mechanisms driving tumor development. GTEx, on the other hand, sourced its samples from 54 non-diseased tissue sites spanning nearly 1000 individuals, providing in-depth perspectives on gene expression dynamics and regulatory patterns across various human tissues. Lastly, TARGET is dedicated to understanding the molecular alterations propelling five pediatric cancers: ALL, AML, kidney cancer, neuroblastoma, and osteosarcoma.

While all three datasets encompass RNA-Seq data, only the CCLE offers distinct measurements for microRNA. For our transportability experiment, we retrieved log2(TPM + 0.001) RNA-Seq data for TCGA, GTEx, and TARGET from the Xena browser [29] and both log2(TPM + 0.001) RNA-Seq data and RPM-normalized microRNA data for the CCLE from the DepMap project [30]. The Xena browser, by providing uniformly realigned and re-called gene and transcript expression data for TCGA, TARGET, and GTEx, assists in minimizing variances in gene-level RNA-Seq data attributable to different bioinformatics pipelines.

Our study focuses on 942 samples from the CCLE which contain measurements for 18,706 mRNAs and 743 microRNAs. The count of RNA-Seq samples from the Xena browser’s TCGA, GTEx, and TARGET datasets [29] stands at 10,013, 7792, and 734, respectively, with all three datasets capturing data for 19,069 genes.

### 2.2. Biological Relevance of Chosen Features

Our feature selection methodology, Thresher (see Section 4.1), first identified 28 miRNA and 28 TF clusters from the TCGA’s FPKM-UQ-normalized dataset. We then selected the medoid of each cluster as a key feature for downstream analysis. Prior studies have shown that the TF [21] and miRNA [22] clusters recognized by the Thresher method are associated with specific biological processes. Consequently, these clusters can be viewed as “regulatory components” representing cell states, as opposed to “principal components”. We employed the unsupervised, nonlinear t-SNE projection technique [45] to the 8895 × 56 input matrix consisting of selected TFs and miRNAs and visualized the outcome in a two-dimensional plot, as shown in Figure 1A. To enhance our comprehension of the patterns, we color-coded the plot based on known cancer types. Overall, the majority of cancer types could be differentiated using the 56 chosen regulatory molecules. Utilizing the TPM-normalized version of the TCGA data yielded comparable results. In this instance, we identified 28 miRNA clusters but observed a reduction in the number of TF clusters to 19. The corresponding t-SNE plot can be found in Supplement Figure S1A,B.

**Figure 1.**
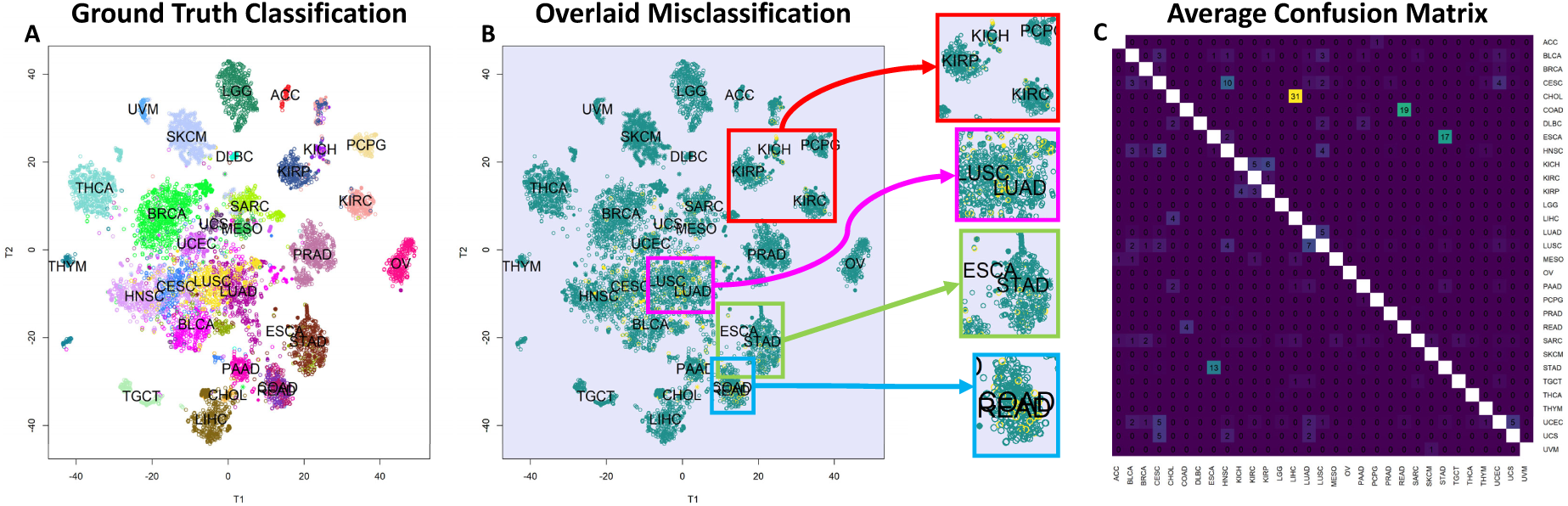
Cancer type classification and visualization through regulatory molecules. (**A**) A 2D t-SNE plot of the 8895 × 56 input TF and miRNA expression matrix, consisting of selected TFs and miRNAs, with colors corresponding to known cancer types, as listed in Supplemental Data, showcasing the ability to differentiate most cancers using 56 regulatory molecules. Note that COAD (Colon Adenocarcinoma) and READ (Rectal Adenocarcinoma) appear almost superimposed due to their high similarity, causing their labels to overlap. (**B**) An enlarged t-SNE plot focusing on misclassified samples, marked in yellow, indicates that classification errors are more common in highly similar cancer types. (**C**) An average confusion matrix generated from 10-fold cross-validation, exposing the error pattern of the SVM classifier with 92.8% accuracy. The numbers within each cell of the heatmap represent the average percentage of times, across 10-fold cross-validation, that samples from one class (row) are predicted to belong to another class (column). A value of 100% would indicate that all samples of a given class were consistently misclassified as another specific class across all cross-validation folds. Diagonal cells representing correct classifications have been left blank for clarity. Lighter matrix entries (excluding diagonals) represent larger errors, and the error pattern predominantly coincides with the t-SNE visualization of errors in panel B. Classifying similar cancer pairs like STAD and ESCA, COAD and READ, and LUSC and LUAD and higher-order similar cancers such as KIRP, KIRC, and KICH, as well as squamous cancers (or cancers with a squamous-cell subtype) including LUSC, CESC, HNSC, and BLCA, is challenging.

### 2.3. High-Accuracy Tissue Classification with 56 Regulatory Molecules

Our analysis (see Section 4.2.1) used 56 regulatory molecules to perform multi-class tissue classification using a vanilla SVM classifier. Figure 1C illustrates the average confusion matrix obtained from 10-fold cross-validation, where the average accuracy of the classifier is 92.8%. However, the accuracy over tissue pairs is bimodal, indicating that the model is able to distinguish most pairs with 100% accuracy but struggles to classify certain pairs. Notably, challenging tissue pairs include COAD (Colon Adenocarcinoma) and READ (Rectal Adenocarcinoma), LUSC (Lung Squamous-Cell Carcinoma) and LUAD (Lung Adenocarcinoma), and ESCA (Esophageal Carcinoma) and STAD (Stomach Adenocarcinoma). Additionally, higher-order classification confusion was observed for KIRP (Kidney Renal Papillary Cell Carcinoma), KIRC (Kidney Renal Clear Cell Carcinoma), and KICH (Kidney Chromophobe), which are all subtypes of renal cell carcinoma, as well as squamous cancers including LUSC, CESC (Cervical Squamous-Cell Carcinoma), HNSC (Head and Neck Squamous-Cell Carcinomas), and BLCA (Bladder Carcinoma, with a basal–squamous subtype). The t-SNE plot of the clusters is shown in Figure 1B, with misclassified samples highlighted in yellow. The plot is zoomed in to emphasize that errors are more frequent in tissues that are highly similar. When using the TPM-normalized data, the classification accuracy slightly decreased to 90.24%, though the misclassification patterns closely mirrored our observations obtained with FPKM-UQ. These results are detailed in Supplement Figure S1C.

### 2.4. Highly Accurate Prediction of Gene Expression Using 56 Selected Features

Our analysis employed the 56 selected features to predict the expression of approximately 20,000 genes using our proposed Tissue-Agnostic and Tissue-Aware models (see Section 4.2.2). Figure 2A,B illustrate the *R*^2^ distribution histogram for both models, revealing that both histograms are mixtures of two components. The tail analysis procedure used beta mixture distributions fitted to the histograms, with the green curve representing the component covering the right tail (i.e., well-explained genes) and the red curve representing the left tail (i.e., poorly explained genes). Notably, the distribution of poorly explained genes is thicker and skewed toward zero in the Tissue-Agnostic model in comparison to the Tissue-Aware model. Additionally, the variance explained by the model is significantly improved in the Tissue-Aware model. The average *R*^2^ values for the Tissue-Agnostic and Tissue-Aware models are 0.45 and 0.70, respectively. When conducting similar experiments with TPM-normalized data, we observed qualitatively similar histograms, presented in Supplement Figure S2A,B. Both histograms for the Tissue-Agnostic and Tissue-Aware models were identified as mixtures of two beta distributions, with average *R*^2^ values of 0.46 and 0.68, respectively.

**Figure 2.**
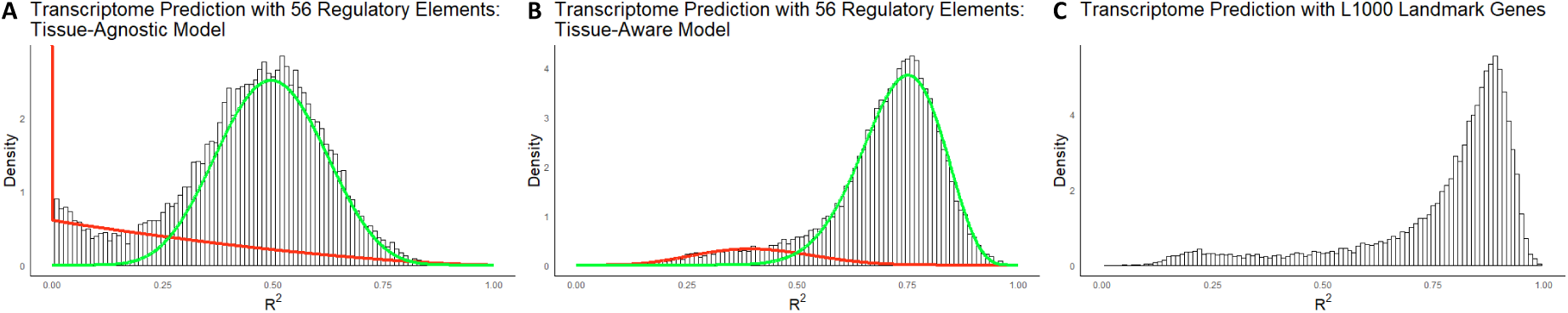
Gene expression prediction and comparison of predictive power between selected regulatory molecules and L1000 genes. (**A**,**B**) *R*^2^ histograms for predicting 20,289 gene expressions using 8895 samples in TCGA with Tissue-Agnostic and Tissue-Aware models, respectively, illustrating the mixture of two components: well- explained genes (green curve) and poorly explained genes (red curve). The Tissue-Aware model demonstrated enhanced performance, with an average *R*^2^ value of 0.70 compared to 0.45 for the Tissue-Agnostic model. (**C**) *R*^2^ distribution of the linear models fitted using 1000 landmark genes [14], displaying an average *R*^2^ value of 0.77. The distribution exhibits a more pronounced bimodality, indicating that, compared to the Tissue-Aware model, some genes are better explained, while the poorly explained tail is also thicker. The Tissue-Aware model achieved comparable predictive power using only 56 predictors according to the *R*^2^ measure and attained a correlation score of 0.9998 compared to 0.9975 for the L1000 genes, highlighting the effectiveness of the 56 selected regulatory molecules in predicting the transcriptome.

In addition, we extracted 345 tissue-enriched genes from the Human Protein Atlas [46] across nine matching TCGA tumor types. These genes exhibit high expression specific to one tissue (e.g., pancreas, ovary, or bladder). We then compared the performance of our Tissue-Agnostic and Tissue-Aware models in predicting the expression levels of these markers within the corresponding TCGA samples. As shown in Figure 3, the Tissue-Aware framework significantly outperforms the Tissue-Agnostic approach on this subset.

**Figure 3.**
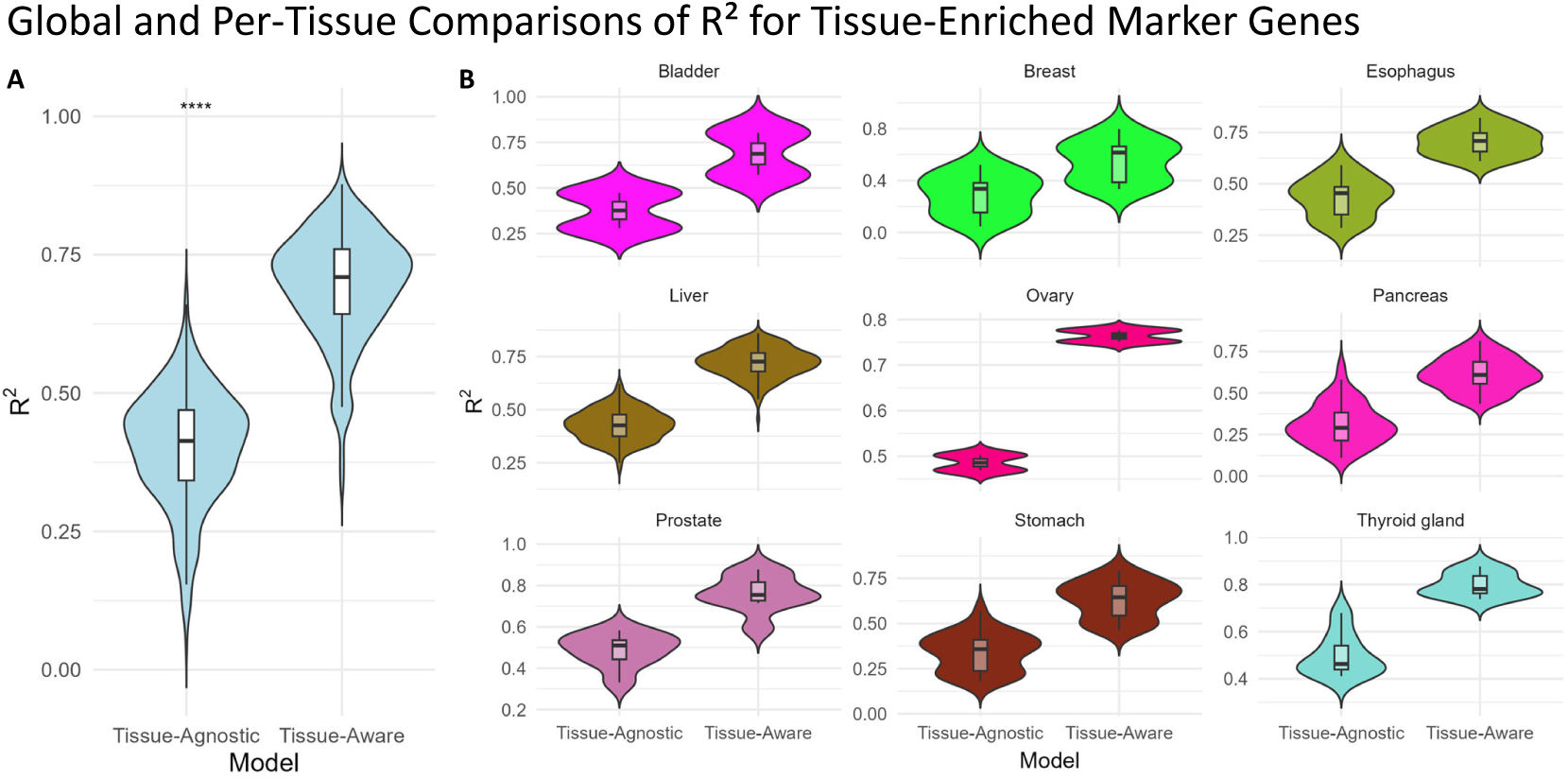
Model performance for tissue-enriched marker genes. We identified 345 marker genes from the Human Protein Atlas that are enriched in 9 TCGA cancer types (i.e., each gene shows high expression in a specific tissue). Because these genes are tightly linked to tissue identity, accurately predicting their expression can be especially important for understanding tissue-specific regulatory mechanisms. Here, we compare our Tissue-Agnostic and Tissue-Aware models on this focused gene set. (**A**) The global violin plot shows a significant overall improvement from Tissue-Agnostic to Tissue-Aware models (paired Wilcoxon test *V* = 0, *p <* 2.2 × 10^−16^, **** denotes statistical significance at *p <* 0.0001).(**B**) A per-tissue breakdown demonstrates that most tissues also show notable improvements when incorporating tissue context.

### 2.5. Comparison of Predictive Power of Selected Regulatory Molecules and L1000 Genes for Transcriptome

In this experiment, we compared the predictive power of our 56 selected regulatory molecules to that of the L1000 genes. Figure 2C shows the *R*^2^ distribution of the linear model fitted using L1000 genes, with an average *R*^2^ value of 0.77 over the entire transcriptome. In contrast, our Tissue-Aware model, using only 56 predictors, has an average *R*^2^ value of 0.70, indicating comparable predictive power. The *R*^2^ distribution of the L1000 genes also exhibits a similar bimodal shape as observed in the Tissue-Aware model. Furthermore, while the L1000 genes reach a correlation score of 0.9975, our Tissue-Aware model achieves a score of 0.9998. These results demonstrate that our 56 selected regulatory molecules are almost as good predictors of the transcriptome as the widely used L1000 genes and, in some cases, even outperform them.

### 2.6. Identifying Pathways Regulated by TFs, miRs, and Alternative Mechanisms

To systematically categorize genes based on their predictability under the Tissue-Agnostic and Tissue-Aware models, we applied both a mixture model classification and a global thresholding approach. The mixture model classified genes into poorly or well-explained categories separately for each model, allowing us to identify genes that switch classes when transitioning from the Tissue- Agnostic to the Tissue-Aware framework.

Additionally, we employed global thresholds of *R*^2^ < 0.3 for poorly explained genes and *R*^2^ > 0.7 for well-explained genes. A gene was categorized as poorly explained only if its *R*^2^ value remained below 0.3 in both models, and as well explained only if its *R*^2^ exceeded 0.7 in both models. Genes that switched categories were identified through the mixture model classification, ensuring that transitions were not solely due to thresholding effects.

Using these criteria, we identified the following:

- 420 genes that remained **poorly explained** across both models.
- 785 genes that remained **well explained** in both models.
- 1107 genes that **switched from poorly to well explained** under the Tissue-Aware model based on the mixture model classification.
- 164 genes that not only switched from poorly to well explained but also exhibited a **4-fold or greater increase** in *R*^2^, representing the strongest improvements in predictability.

We visualize these results in Figure 4, where Panel A presents a direct comparison of gene predictability (*R*^2^) across models. Genes that remained poorly explained are shown in red, consistently well-explained genes in green, and genes that transitioned from poorly to well explained in blue. Additionally, those that demonstrated at least a 4-fold increase in predictability are highlighted in orange.

**Figure 4.**
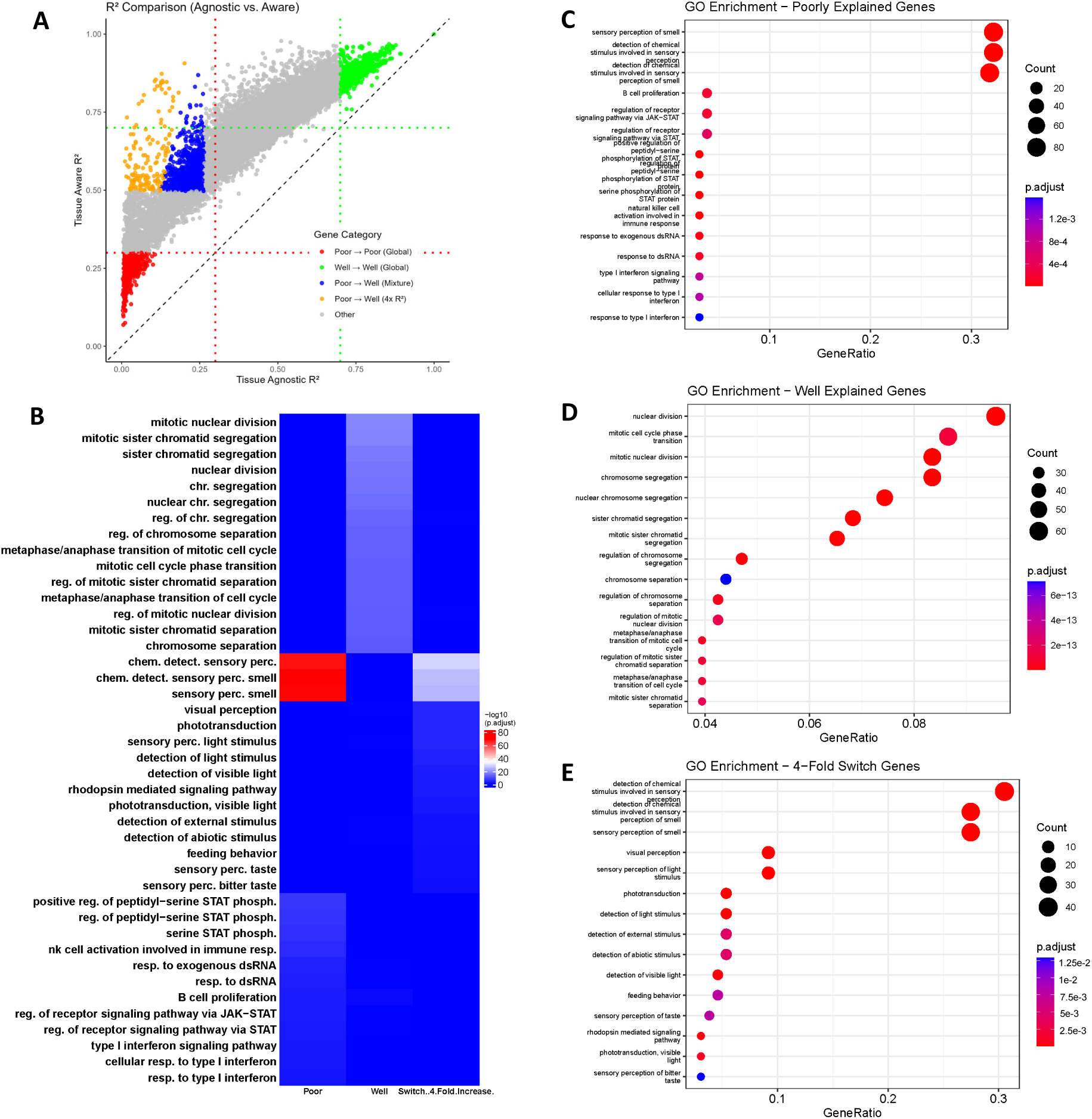
Comparative analysis of gene predictability and enrichment pathways. (**A**) Scatter plot comparing gene predictability (*R*^2^) in the Tissue-Agnostic and Tissue-Aware models. Genes that remain poorly explained in both models are shown in red, consistently well-explained genes in green, and genes that transition from poorly to well explained in blue. Genes that transition from poorly to well explained and exhibit at least a 4-fold increase in predictability are highlighted in orange. Notably, all points lie above the diagonal, indicating that all genes’ predictability increases when moving to the Tissue-Aware model. (**B**) Heatmap of pathway enrichment significance (− log_10_(p.adjust)) across gene groups. All three categories of genes show distinct enrichment patterns, while some pathways related to sensory perception display partial overlap between switch and poorly explained genes. (**C**) Gene Ontology (GO) enrichment dot plot for genes classified as poorly explained in both models. Top enriched pathways include immune signaling and sensory perception functions. (**D**) GO enrichment dot plot for genes classified as well explained in both models, highlighting pathways involved in cell cycle regulation and mitosis. (**E**) GO enrichment dot plot for genes that transitioned from poorly to well explained. Enriched pathways are primarily related to sensory perception and environmental stimulus response.

To further characterize the biological roles of these gene groups, we performed Gene Ontology (**GO**) enrichment analysis, identifying significantly enriched biological pathways in each group (Figure 4):

- Poorly explained genes (Panel C) were predominantly associated with *immune signaling and sensory perception* pathways, such as the *type I interferon response*, *STAT phosphorylation*, and *B-cell proliferation*.
- Well-explained genes (Panel D) were strongly enriched for *cell cycle and mitotic processes*, including *chromosome segregation*, *nuclear division*, and *sister chromatid separation*.
- Genes that switched from poorly to well explained (Panel E) were significantly enriched in pathways related to *sensory perception and stimulus response*, particularly *phototransduction*, *detection of light stimuli*, and *feeding behavior*.

Finally, we summarized the pathway-level similarities and differences across gene groups using a heatmap (Figure 4B), where pathway enrichment significance is represented by − log_10_(p.adjust). The heatmap presents the top pathways enriched in each gene category: poorly explained genes (Poor), well-explained genes (Well), and genes that switched from poor to well explained with a four-fold increase in *R*^2^ (Switch). The pathways are arranged in rows, while the three gene categories are displayed in columns. The color intensity represents the statistical significance of pathway enrichment, with red indicating highly significant pathways and blue indicating non-significant or weak enrichment.

### 2.7. Distributional Shift and Mismatch of microRNAs Compromise Tissue-Aware Model’s Transportability

So far, we have used the tissue of origin from TCGA in the Tissue-Aware model, but tissue labels do not always align across datasets, making direct matching challenging. To address this, we introduced Pseudo-Tissues, where samples with similar gene expression profiles are clustered within the source dataset. In the target dataset, samples are assigned to Pseudo-Tissues based on their proximity to the centroid of each cluster (see Section 4.6).

To ensure the robustness and reliability of our Tissue-Aware model in predicting gene expression across different datasets, we investigated its generalization error [47] as a function of the number of Pseudo-Tissues. This error metric provides insights into the model’s ability to capture underlying biological patterns in new, unseen datasets without merely fitting to known data. Figure 5A,B depict the cross-validation curves to tune the number of Pseudo-Tissues (clusters) for the Root Mean Square Error (RMSE) and correlation score, respectively, using the FPKM-UQ normalized TCGA dataset. The optimal number of Pseudo-Tissues was found to be 15, based on both criteria. Additionally, we obtained analogous curves with the TPM-normalized data, as shown in Supplement Figure S3A,B, which concur that 15 is the optimal number of Pseudo-Tissues.

**Figure 5.**
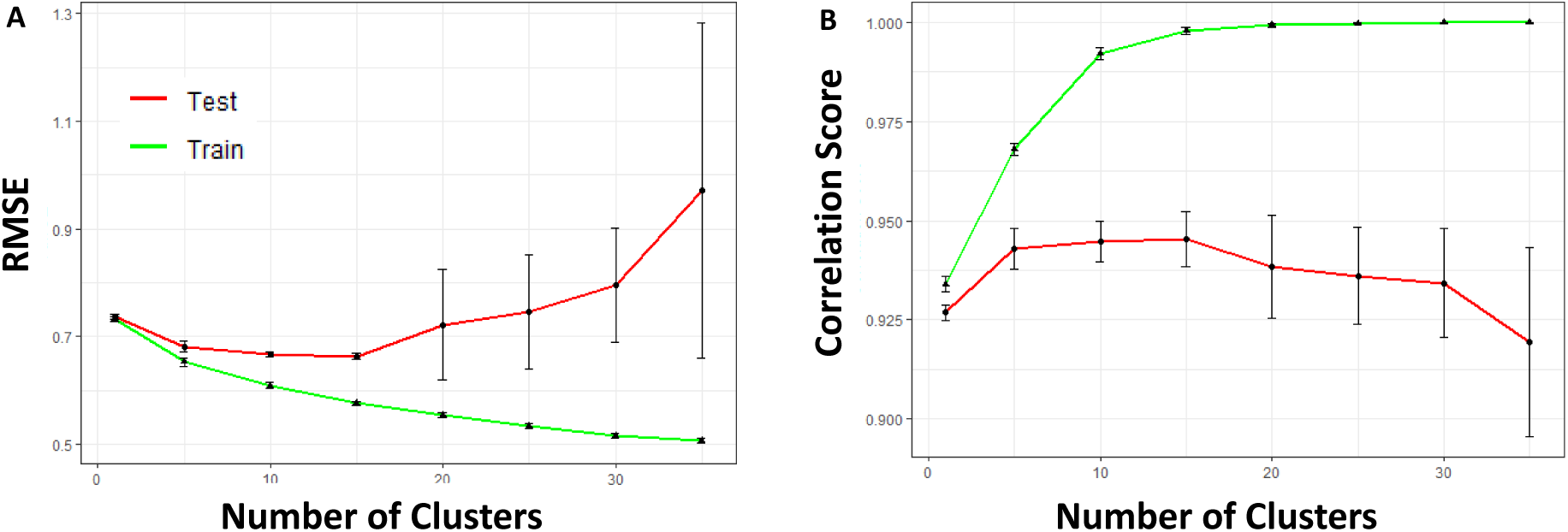
Optimizing Pseudo-Tissues for Tissue-Aware model generalization. (**A**,**B**) Cross-validation curves for tuning the number of Pseudo-Tissues (sample clusters) based on RMSE and correlation score, respectively, using the TCGA dataset. The optimal number of Pseudo-Tissues is determined to be 15 for both criteria. However, cross-validation assumes that the training and testing data share the same distributions, which is not the case when distribution shifts are present, such as when selected miRNA expressions are heavily shifted in the CCLE, thus hindering the transportability of the predictive model.

Nevertheless, the transportability assessment (out-of-distribution generalization [48]) of the model on the CCLE dataset yielded a correlation score of 0.22 and RMSE of 3.83, which were worse than anticipated. We hypothesized that this diminished performance was due to discrepancies in selected miRNAs and their distributional shifts [49] between TCGA and CCLE. One major challenge in model transportability arises from inconsistencies in miRNA measurements across datasets. Unlike TCGA, which provides directional measurements for mature miRNAs (e.g., hsa-miR-21a-5p vs. hsa-miR-21a- 3p), CCLE reports only aggregate counts, leading to mismatches. To investigate the impact of these differences, we quantified the distributional shift of matched features between TCGA and CCLE using the Wasserstein distance [50], as shown in Figure 6. Specifically, the distance was calculated for each matched pair of features (either TF or miRNA) For instance, the density for the 28 miRNAs stemmed from 28 paired TCGA and CCLE miRNA samples. This was then contrasted against a null distribution created from non-matching feature pairs. The findings, as illustrated in Figure 6A,B, revealed that, while matched TFs in the datasets were more aligned than random pairs, the distances for matched miRNAs were comparable to those of random miRNA pairs. Such observations underscore the distinct distributional nature of the matched miRNAs, reinforcing the notion that a model trained on TCGA might encounter challenges when tested on the CCLE.

**Figure 6.**
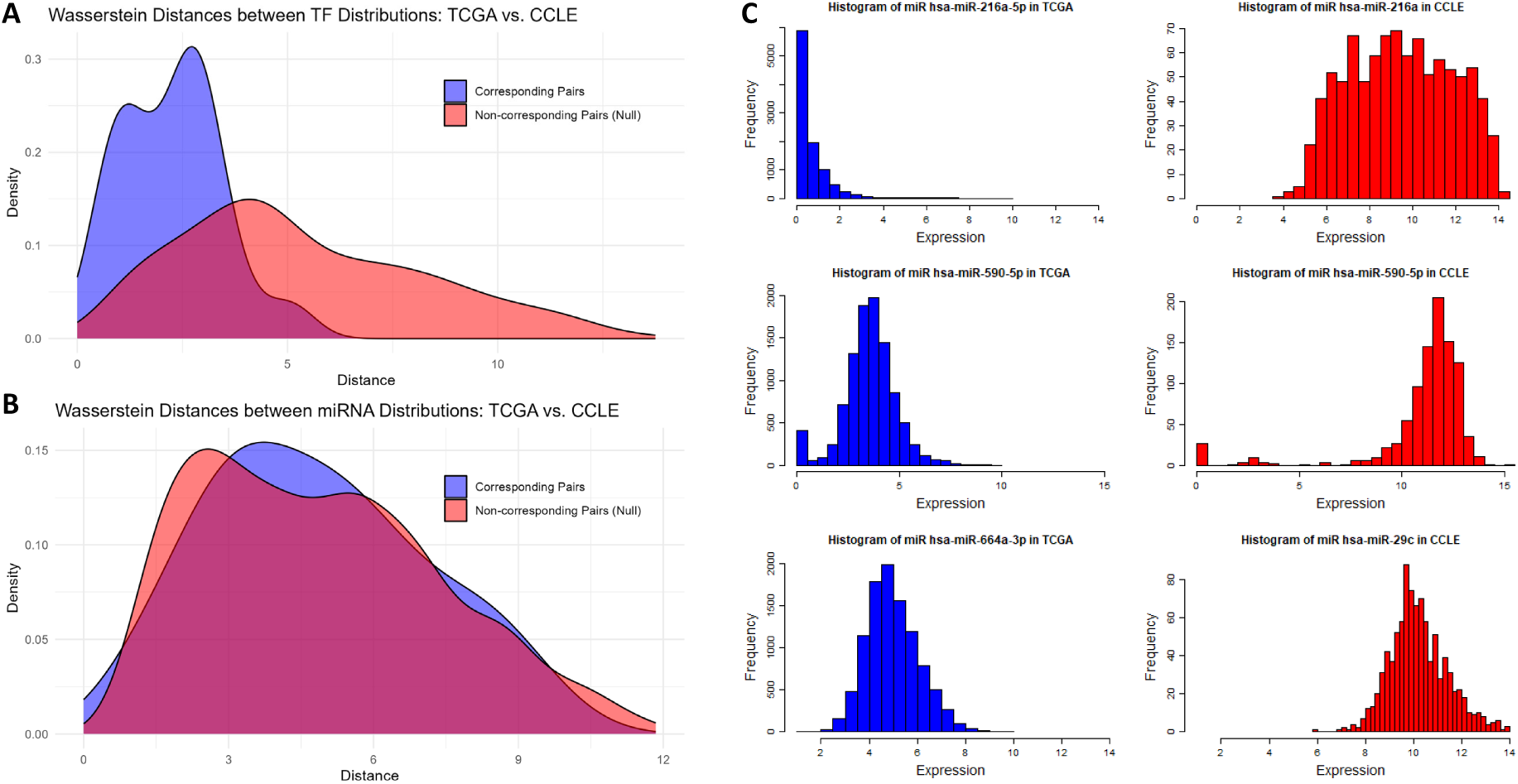
Shift in distributions of regulatory molecules across TCGA and the CCLE affecting model trans- portability. The Tissue-Aware model’s transportability challenges, when incorporating both TFs and miRNAs, contrast its efficacy with TFs alone, pointing toward differential miRNA distributions between TCGA and the CCLE. The visualizations below substantiate this observation. (**A**) Comparing TF distributions: The blue density visualizes the Wasserstein distance between corresponding (matched) TFs from TCGA and the CCLE. It is shifted leftwards relative to the empirical null (red density). The null model is derived using distances from mismatched TF pairs across datasets. The data suggest TF distributions are predominantly congruent between TCGA and the CCLE. (**B**) Comparing miRNA distributions: For miRNAs, the landscape differs. The blue density, signifying the Wasserstein distance for matched miRNAs, aligns with the null (red density), indicating distinct miRNA distributions between the two datasets. (**C**) Dissecting cross-dataset microRNA expression variations: Each row showcases a pair, with blue histograms representing TCGA and red denoting the CCLE. The directionality of miRNAs in TCGA’s may not be captured in the CCLE’s, highlighted by hsa-miR-21a-5p (TCGA) versus hsa- miR-21a (CCLE) in row 1. Although certain miRNAs have their directionality recorded in both datasets, their distribution can still be different due to various biological or technological factors, illustrated by hsa-miR-590-5p in row 2. In five cases, we could not find the miRNAs of TCGA in the CCLE. Our method then opted for the closest alternative, as shown by hsa-miR-664a-3p (TCGA) being paired with hsa-miR-29c (CCLE) in row 3.

The distance distribution offers a broad perspective on the discrepancies between the features of the two datasets. To further delve into individual variability of the features, we illustrate the distributions for three paired miRNAs in Figure 6C. Each of these pairs exemplifies a distinct miRNA matching challenge, as detailed in Section 4. The unavailability of specific mature directional miRNA data in the CCLE introduced potential mismatches, which can be considered as a technological source of covariate shift. This is shown by the histograms for hsa-miR-21a-5p (TCGA) versus hsa-miR- 21a (CCLE) showcased in the top row of Figure 6C. Additionally, among the directionally matched miRNAs, certain ones displayed pronounced distributional shifts, indicative of a biological source of covariate shift. This is illustrated by hsa-miR-590-5p in the second row of Figure 6C. Moreover, in the absence of five crucial miRNAs in the CCLE, they were paired with the most closely related alternatives, encapsulating both technological and biological sources of shift. This is represented in the third row of Figure 6C by hsa-miR-664a-3p (TCGA) being paired with hsa-miR-29c (CCLE). Conversely, the TF distributions exhibited remarkable consistency across the two datasets.

### 2.8. Robust Transportability of Tissue-Aware Model Using Transcription Factors Alone

Given the challenges with miRNA consistency we identified in our initial transportability experiments, we shifted our focus solely toward TFs. This decision was supported by the more stable expression distributions of TFs observed across datasets, as evident in Figure 6A. Utilizing 19 selected TFs from the TPM-normalized TCGA dataset, we trained a Tissue-Aware model and subsequently evaluated its ability to predict the entire transcriptome on the CCLE, GTEx, and TARGET datasets. Remarkably, the model showcased superior performance on the CCLE when relying solely on TFs. The results were an average RMSE of 2.38 and a correlation score of 0.49. This was a notable improvement from the results obtained when both TFs and miRNAs were used as features.

Given the absence of separate miRNA measurements in the GTEx and TARGET datasets, we limited our model to TFs as predictors. The results were promising: For GTEx, we attained an average RMSE of 1.89 and a correlation score of 0.68. For TARGET, the corresponding values were an average RMSE of 2.34 and a correlation score of 0.53. These outcomes across the three distinct datasets underscore the transportability of our Tissue-Aware model. When relying exclusively on transcription factors, the model capably estimated the expression of approximately half of the genes across all target datasets.

## 3. Discussion

### 3.1. The Role of Causal Predictors in Robust Transcriptome Representation

Understanding gene regulatory networks, capturing cell states through gene expression profiles, and obtaining low-dimensional representations of cell states are three essential and interconnected biological challenges that, collectively, offer a cost-effective and robust method to evaluate the impact of various perturbations, encompassing mutations and therapeutic interventions. .

In this study, we demonstrated that, by using our prior knowledge of the critical roles of transcription factors and microRNAs in gene regulation, we can establish a low-dimensional representation of cell states and infer the entire transcriptome from a limited number of regulatory molecules. The value of a reduced cell state representation lies in its ability to capture the gene expression distribution not only under normal conditions but also under various perturbations, such as drugs, mutations, or gene knockouts. These perturbations fundamentally change the joint distributions of gene expressions, generating interventional distributions [51] over expressions that can differ significantly from the normal state. To obtain the minimum number of genes needed to explain most transcriptome variations, it is necessary to understand or infer the gene regulatory network and select master regulators to fully predict the outcomes of perturbations. We hypothesized that using TFs and miRNAs as a shortcut can work in place of full gene regulation inference because of their significant role in the regulation of various biological processes. Consequently, under any perturbation or environment, their values serve as robust predictors of the entire transcriptome. Our hypothesis is supported by two key findings. Firstly, our feature selection pipeline identified regulatory molecule clusters that have biological significance, as reported in [21,22]. Secondly, the selected features demonstrate strong predictive capability in supervised learning tasks across different environments, such as different tissues, and under different perturbations, such as DNA aberrations in cancer cells. Together, these results provide further evidence in support of our hypothesis.

### 3.2. Challenges in Cancer Type Classification

Our findings demonstrate that these selected features could effectively distinguish between 31 cancer types, as illustrated by the t-SNE plot (Figure 1A). Moreover, the classification accuracy obtained by our vanilla SVM classifier was promising at 92.8%, although the model faced some challenges with specific tissue pairs and subtypes. Biologically, the misclassification of these tissues can be rationalized. COAD and READ are very similar subtypes of colorectal cancer [52], while LUSC and LUAD are subtypes of lung cancer and ESCA and STAD are both types of gastrointestinal cancers. A similar situation arises for KIRP, KIRC, and KICH, where all are subtypes of renal cell carcinoma that originate from different cell types in close proximity to the kidney.

### 3.3. Enhanced Gene Expression Prediction Using Selected Features and Tissue-Aware Models

The prediction of gene expression using the 56 selected features was highly accurate, with an average *R*^2^ value of 0.45 for the Tissue-Agnostic model and 0.70 for the Tissue-Aware model. The Tissue-Aware model showed significant improvement over the Tissue-Agnostic model, suggesting that incorporating tissue-specific information into our multi-task learning framework can enhance gene expression prediction. There are two potential biological explanations for the effectiveness of the Tissue-Aware model. First, several other types of regulatory molecules, such as epigenetic modifiers [53] and chromatin remodeling complexes [54], play essential roles in gene regulation as well as TFs and miRs. Some of these, like DNA methylation and histone modification, are crucial in determining cell identity and, ultimately, tissue type [55,56]. Therefore, using tissue type can serve as a surrogate for these omitted regulators. Secondly, from a causal perspective, the interaction between regulatory molecules can cause the causal effect of TFs and miRNAs to differ in each environment. Consequently, our multi-task learning framework, which fits a model per environment with shared parameters across environments, is a more biologically plausible model.

We hypothesized that the causal relevance of TFs and miRs, along with our flexible multi-task learning framework, would enable a succinct representation of the transcriptome. We confirmed this hypothesis by demonstrating that our 56 selected features are as predictive as the widely used L1000 genes. The predictive power of our 56 regulatory molecules was nearly as good as the L1000 genes according to both *R*^2^ and the correlation score measures. Our results highlight the effectiveness of our feature selection method, which is guided by biological knowledge about the causal role of regulatory molecules in transcription.

### 3.4. Pipeline Robustness to Diverse RNA-Seq Normalization Methods

In our analysis, we employed two well-established and commonly used normalization methods: FPKM and TPM. All steps of our study, with the exception of the transportability experiment, were conducted utilizing data normalized by both techniques. The primary distinction observed was in feature selection. Specifically, Thresher, our feature selection method, identified 19 TF clusters with TPM normalization, in contrast to the 28 identified using FPKM. However, the number of determined miRNA clusters remained consistent at 28 for both methods. While there was a 16% reduction in the total number of selected features ((28-19)/56) when employing TPM instead of FPKM, this reduction did not result in a significant performance drop in tissue type classification or transcriptome prediction. These observations underscore the robustness of our pipeline, irrespective of the chosen normalization method.

### 3.4. Biological Interpretation of Gene Enrichment Results

#### 3.5.1. Immune and Sensory Pathways Dominate the “Poorly Explained” Category

As shown in Figure 4C, genes that remained poorly predicted in both the Tissue-Agnostic and Tissue-Aware models were significantly enriched in immune signaling (e.g., type I interferon response, STAT phosphorylation) and sensory perception. In the case of immune-related genes, their expression levels can shift dramatically in response to infection, inflammation, or other external cues, and they often exhibit high heterogeneity across individuals and tissues [57,58]. Such variability can exceed what our limited panel of 28 microRNAs and 28 TFs can capture.

For the sensory perception category, a large fraction of the enrichment signal stems from olfactory (smell) genes. Because the TCGA dataset comprises primarily cancer samples, we did not have dedicated nasal or olfactory tissues available for analysis. As a result, the Tissue-Aware model still lacks the specialized context required to capture these genes’ regulation. Nonetheless, when tissue context was included, we could explain many other stimulus-driven pathways (e.g., light detection, feeding behavior), which we discuss further in Section 3.5.4. This underscores the importance of precise tissue information in modeling genes that respond to environmental or physiological stimuli, although the absence of olfactory tissue data continues to limit our ability to account for smell-associated processes.

#### 3.5.2. Cell Cycle and Mitosis Genes Are Consistently “Well Explained”

In contrast to the poorly explained categories, the genes that remained well explained under both models (Figure 4D) were strongly enriched for fundamental processes such as chromosome segregation, nuclear division, and sister chromatid separation [59–61]. These core processes appear to be governed by robust TF and miRNA networks (e.g., the E2F family, MYC), whose regulatory influence is relatively consistent across different tissue types.

Because many of these genes function as “housekeeping” elements that drive essential cellular functions, they exhibit a level of tissue-independent expression [62]. This stability makes them easier to capture with the fixed panel of TFs and miRNAs, thereby explaining their higher predictability in both the Tissue-Agnostic and Tissue-Aware frameworks. Collectively, this finding underscores that critical cell cycle regulators are less sensitive to context-specific variation, and hence are more amenable to accurate computational modeling.

#### 3.5.3. Gains in Predictability Reveal Tissue-Dependent “Switch” Genes

A set of 1107 genes transitioned from “poorly explained” in the Tissue-Agnostic framework to “well explained” once tissue information was integrated (Figure 4A). This suggests that the Tissue-Aware model can better capture the context-dependent interplay between these genes and their regulatory factors and shows how local regulatory environments and specialized signals shape gene expression.

#### 3.5.4. The Strongest Improvements: 4-Fold Increase in Predictability

Within the “switch” group, 164 genes showed at least a 4-fold increase in *R*^2^. The gene enrichment analysis (Figure 4A,B,E) suggested that many of these genes lie in sensory perception pathways, including the detection of chemical stimuli, smell, taste, and vision. Their large gains under Tissue-Aware modeling suggest that these processes are highly dependent on local tissue context or specialized cell types. These observations further support the notion that incorporating tissue-level factors is vital for accurately characterizing genes that respond to external signals or exhibit strong context dependence.

### 3.6. Factors Limiting the Prediction of Poorly Predicted Genes

Although our Tissue-Aware model significantly improved transcriptome-wide prediction accuracy, certain genes remain challenging to predict accurately. Here, we examine several potential factors that may explain these limitations.

#### 3.6.1. Alternative Regulation in Poorly Explained Genes

Genes poorly explained by both Tissue-Agnostic and -Aware models may not be heavily regulated by TFs and miRs. These genes might be regulated by other mechanisms, such as signaling pathways [63], post-translational modifications [64], protein–protein interactions [65], and epigenetic regulation [55]. These alternative mechanisms may allow rapid and dynamic control of cellular responses, immune defense, and maintenance of cellular integrity.

#### 3.6.2. Omission of Tissue-Specific microRNAs

Another plausible explanation for our model’s difficulty in accounting for certain poorly explained genes stems from our exclusion of microRNAs that have zero expression in 75% of the samples. Given that the expression and function of some miRNAs can be tissue specific, it is possible that those we omitted might serve pivotal roles in regulating genes within those tissues. Thus, omitting them could hinder the model’s capacity to adequately explain some genes.

#### 3.6.3. Low Transcriptomic Presence of Poorly Predicted Genes

Further analysis revealed that the average expression of the genes that our model struggled to predict consistently ranked in the bottom 6% when compared to the average expression of all genes. Due to their context-specific, low-level expression, our model may lack the sensitivity required for accurate prediction of these genes.

### 3.7. Impact of Covariate Shift on Transportability

The transportability of the Tissue-Aware model was tested on the CCLE dataset. Despite the promising performance on the TCGA dataset, covariate shift and mismatches, especially in microRNA expressions, affected the model’s prediction accuracy on the CCLE dataset. Both the global picture of the shift in miRNAs as depicted in Figure 6B and the individual examples illustrated in Figure 6C support the idea that the shift in miRNA distributions is significant and that they are a culprit in the lack of transportability of our model. By focusing solely on TFs in our final experiment, the model’s transportability performance significantly improved, confirming the detrimental effects of including mismatched or shifted miRNAs in the predictions. This observation emphasizes the necessity of ensuring the robustness of features across diverse datasets to improve model transportability. The pronounced discrepancies in microRNA expressions between the datasets could be attributed to variations in experimental methodologies, preprocessing steps, or inherent biological differences.

### 3.8. Significance of Improved Transportability with the TF-Only Tissue-Aware Model

The marked improvement in transportability achieved when solely employing TFs for predictions underscores the importance of transcription factors. As primary regulators of gene expression, they appear to offer a more stable and consistent feature set across diverse datasets. Yet it is worth noting that the transportability performance did not match the training and testing outcomes from the TCGA datasets, suggesting opportunities for further refinement. Numerous potential research directions which could enhance transportability present themselves. These include implementing domain adaptation techniques [49] to address covariate shifts in TFs or tapping into methods from the multi-output prediction literature [66], where harnessing the inherent correlations between outcomes, here, the predicted gene expressions, is a tangible opportunity to elevate overall prediction accuracy.

## 4. Materials and Methods

This section describes the methodology used in this study, including feature selection, predictive modeling, and model evaluation across datasets. A detailed description of the datasets and preprocessing steps is provided in Section 2.1.1. Our methods aim to identify a compact set of regulatory molecules—transcription factors (TFs) and microRNAs (miRNAs)—that effectively capture transcriptomic complexity while ensuring generalizability across different tissue types and datasets.

### 4.1. Feature Selection

In this study, we aim to understand the computational complexity of transcriptomes by selecting a low-dimensional representation that captures most of the information about the sample’s state. Specifically, we focus on regulatory molecules miRNAs and TFs as features and aim to identify a small subset of them that can be used in downstream predictive analyses while maintaining comparable performance to using the whole transcriptome.

We chose feature selection instead of dimensionality reduction methods like Principal Component Analysis (PCA) because measuring the corresponding low-dimensional latent features, i.e., linear combinations of all TFs and miRNAs, requires measuring all regulatory molecules, which can be expensive and challenging for low-quality RNA samples. Additionally, interpreting the biological relevance of latent features that are superpositions of numerous genes can be challenging [31]. We chose not to use prediction-guided feature selection methods [32], such as LASSO [33], as our goal was to choose features in a task-agnostic manner, ensuring that the performance of any downstream prediction task remains comparable to the situation where the entire transcriptome is measured. For example, employing LASSO for whole-transcriptome prediction could yield varying sets of predictive miRNAs and TFs for each gene, making the measurement of nearly all miRNAs and TFs a necessity. This goes against our aim of showing that a subset of regulatory molecules can efficiently represent the transcriptome.

Our feature selection process employs the Thresher package [34], which automatically selects the optimal number of clusters for clustering TFs and miRNAs into regulatory components, shown in Figure 7. We then select the medoid of each cluster as the representative gene for the corresponding regulatory component. The Thresher pipeline begins by applying PCA to the feature-by-sample matrix (instead of the sample-by-feature matrix) to derive a low-dimensional representation of each feature. To identify the number of significant principal components, we use a variant of the Bayesian model selection method [35], implemented in the PCDimension package. We further refine the feature set by removing outliers using a length threshold for each feature representation. A threshold of 0.35 was chosen based on several simulation studies [34]. Finally, we cluster the *directions* of feature representation vectors, corresponding to points on the hypersphere, using the von Mises– Fisher mixture model [36]. We select the optimal number of regulatory molecule clusters using the Bayesian Information Criterion [37]. To demonstrate that the selected regulatory molecules capture most of the transcriptome’s information, we perform various predictive tasks, summarized in the subsequent sections.

**Figure 7.**
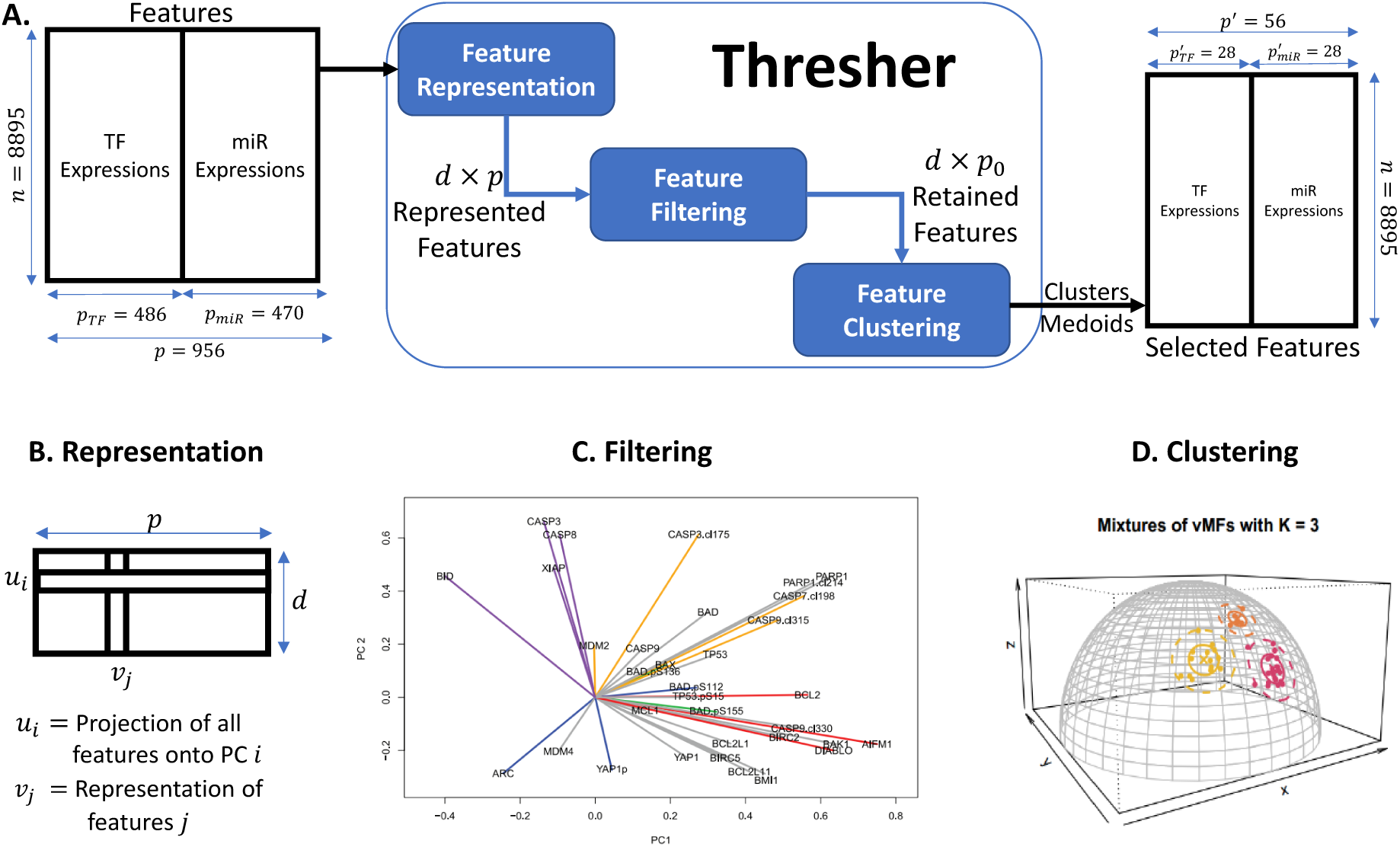
Overview of the feature selection pipeline. (**A**) Overview of Thresher, detailing its input, internal processes, and output: The regulatory molecules of the TCGA dataset, i.e., transcription factors and microRNAs, serve as the input. Thresher first learns a representation of each TF and microRNA, then filters out the less significant ones and finally clusters the remaining miRs and TFs. A medoid from each cluster is chosen to represent it. The resulting selected regulatory molecules are two orders of magnitude fewer than the initial set but retain comparable biological predictive power. Panels (B–D) provide detailed illustrations of each internal step of the Thresher pipeline. (**B**) Feature representation: Features are represented using PCA applied to the feature-by-sample matrix. Each feature *i* is represented by *ν_i_* ∈ R*^d^*, where *d* is determined through Bayesian model selection. (**C**) Feature filtering: Feature representations are visualized using only their first two principal components, revealing significant variance in the lengths of loadings. Features with smaller loadings are filtered out. Specifically, a feature is discarded if ∥*v_i_* ∥_2_> 0.35. (**D**) Feature clustering: Post-filtering, the lengths of feature representations are considered less critical. The clustering then focuses on the directions of the features, *v*_*i*_ /*v*_*i*_ ∥_2_, which requires the clustering of points on a hypersphere, where features are assumed to have mixtures of von Mises–Fisher distributions.

### 4.2. Assessing Predictive Power of Selected Regulatory Molecules

#### 4.2.1. Tissue Classification Using Selected Regulatory Molecules

To show that the 56 selected regulatory molecules are biologically informative, we used them to predict the cancer type of TCGA samples. This was a multi-class classification problem [38] with 31 outcome labels. To train our classifier, we used a Support Vector Machine (SVM) algorithm [39] and report the average generalization error on hold-out data through 10-fold cross-validation.

#### 4.2.2. Transcriptome Prediction Using Selected Features and Tissue Type Information

Our goal was to test whether the 56 selected features provided an accurate representation of the entire transcriptome. In the first experiment, we used linear regression to predict the entire transcriptome using the selected features. We had approximately 20,000 regression tasks, with each task consisting of 56 features and ∼9000 samples. For each outcome gene, we obtained a model with 57 parameters, consisting of 56 coefficients and one intercept. Since we do not use cancer tissue type information in prediction, we call this the **Tissue-Agnostic** model.

In the second experiment, we used the cancer tissue type information available in the TCGA dataset to improve our model and call it the **Tissue-Aware** model. Instead of predicting the outcome for each gene as a single task, we predicted the outcome in each tissue as a separate task, resulting in 31 tasks for TCGA. We first fitted a global regression model across all tissues and then fitted the residuals of the global model in each tissue through a secondary regression. This can be considered a form of multi-task learning [40], where a shared intercept is learned for all tasks along with one linear model per task [22,41,42].

Mathematically, to predict the expression of gene *g*, denoted as **y***^g^* ∈ R*^n^*, in tissue *t* using the selected expression matrix **X** ^56^ and an *_t_* of TFs and miRs, we determined a coefficient vector *β_t_* ∈ R intercept *α_t_* specific to each tissue. Additionally, we introduced a global (cross-tissue) intercept *α*_0_. This Tissue-Aware optimization is expressed as

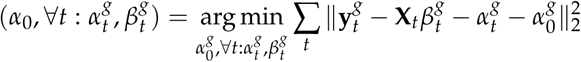

### 4.3. Biological Insights from Evaluating Transcriptome Prediction Accuracy

To gain insights into the biological implications of transcriptome prediction accuracy, we evaluated the Tissue-Agnostic and Tissue-Aware models and examined how much variance in the transcriptome they accounted for. Our analysis focused on the distribution of *R*^2^ values, which represents the proportion of variation in gene expression that each model explains. We created a histogram of *R*^2^ values for the nearly 20,000 genes predicted and fitted a beta mixture model to identify poorly, moderately, and well-explained genes from the histogram. We applied the expectation–maximization algorithm with a range of mixture components (*k* = 1, 2, 3, 4) and used the Bayesian Information Criterion (BIC) score to determine the optimal number of components. The *R*^2^ distribution of both models showed evidence of two beta distribution components, which we labeled as **poorly and well-explained components**, shown in Figure 2.

Additionally, we sought to characterize the functional pathways associated with genes whose predictability differs between the two models. First, we applied global *R*^2^ thresholds to define *poorly explained* genes (*R*^2^ < 0.3 in both models) and *well-explained* genes (*R*^2^ > 0.7 in both models). We then identified a set of *“switch” genes* that change from poorly to well explained when moving from the Tissue-Agnostic to the Tissue-Aware model, as determined by mixture modeling, and highlighted a subset of these genes exhibiting at least a four-fold increase in *R*^2^. For each of these categories (global poorly explained, global well explained, and switched/4-fold improved), we performed Gene Ontology (GO) enrichment analysis (using ToppGene [43]) to uncover pathways that are either strongly driven by the interplay of TFs and miRNAs or remain difficult to predict under our current approach.

### 4.4. Evaluation of Normalization Method Influence on Results

Throughout our study, we primarily utilized the FPKM-UQ normalization method for TCGA’s RNA-Seq data. To ensure our results are not solely dependent on this specific normalization method, we conducted all experiments—including tissue classification, transcriptome prediction, and gene enrichment analysis—using TPM-normalized TCGA data. TPM is recognized for better preservation of biological signals compared to other normalization techniques [44].

### 4.5. Comparison of Performance Between Selected Regulatory Molecules and L1000 Genes

The L1000 Gene Expression Signature Profiling Technology and the performance of the 978 landmark genes it uses for whole-transcriptome prediction have been widely studied to identify new therapeutic targets and drug combinations [14]. However, due to the lack of access to the dataset used for their gene selection and prediction, we could not test the performance of our 56 selected regulatory molecules on the same dataset. To ensure a fair comparison, we instead used the L1000 genes to fit the TCGA data from scratch.

We compared our Tissue-Aware model, which employs 56 genes, and the linear model, which uses L1000 genes. To evaluate the performance of the models, we computed their average *R*^2^ scores and correlation scores. The correlation score, introduced in the L1000 landmark paper [14], measures the algorithm’s overall performance in predicting gene expression as the fraction of genes that are “best inferred”. A gene *g* is best inferred if the correlation *R_gg_* = cor(**y**^ *^g^*, **y***^g^*) is significant. To determine if *R_gg_* is significant, a null distribution of correlation is made by randomly pairing genes *g*_1_ and *g*_2_ and computing *R_g_*_1_ *_g_*_2_ = cor(**y**^ *^g^*^1^, **y***^g^*^2^). The inference is deemed accurate if *R_gg_* is higher than the 95th percentile of the null distribution. The linear model trained on the data from [14] that uses L1000 landmark genes as features has a reported correlation score of 81%.

### 4.6. Assessing the Model’s Transportability Across Datasets

To validate the transportability of our Tissue-Aware transcriptome prediction model, we applied it to an independent dataset: the CCLE (Cancer Cell Line Encyclopedia) project [28]. The predictors for mRNA expression were sourced from the TFs and miRNAs selected by Thresher using TCGA data, as shown in Figure 7.

However, integrating the selected miRNAs from TCGA with the CCLE posed several challenges. Notably, TCGA includes measurements for both mature miRNA directions, such as hsa-miR-145-3p and hsa-miR-145-5p, whereas the CCLE only reports hsa-miR-145. Given the likely disparities in the abundance and targets of the 3p and 5p mature miRNAs, accurate matching to the CCLE’s miRNAs is crucial. Yet it remains unclear if the CCLE captures a dominant variant or an aggregate count of both. We opted to match the directional miRNAs from TCGA to their non-directional counterparts in the CCLE, understanding that, depending on the CCLE’s measurement methodology, this could significantly impact prediction performance. Furthermore, five miRNAs selected from TCGA are absent in the CCLE. For these, we substituted the second “closest to the centroid” member from the respective miRNA clusters identified by Thresher. An additional concern arose from observing notable distributional differences for some matched miRNAs between TCGA and the CCLE. We explore these topics further in subsequent sections.

With the Tissue-Aware model, our aim was to predict the transcription of 18,706 mRNAs across 942 CCLE samples. This task necessitated establishing a correspondence between the tissue (or cancer type) labels of TCGA and the CCLE. Instead, we clustered TCGA samples with similar gene expressions into “Pseudo-Tissues”. We employed the K-means algorithm [37], treated the number of clusters *K* as a hyper-parameter, and determined its optimal value via 5-fold cross-validation. To evaluate model performance on the CCLE, each sample was mapped to a TCGA Pseudo-Tissue using the minimum Euclidean distance. The model trained on the associated Pseudo-Tissue was then deployed to predict the complete transcriptome of the given CCLE sample.

### 4.7. TF-Based Model Transportability Across Datasets

Considering the frequent omission of miRNAs in many large-scale RNA-seq datasets, we explored the feasibility of exclusively using TFs for whole-transcriptome prediction. This shift towards TFs not only simplifies the prediction model but also widens the scope of applicable RNA-Seq datasets for our experiments. While we could potentially have utilized any RNA-Seq dataset, we favored those with broad tissue representation, ensuring the robustness of our predictions across both cancerous and non-cancerous contexts.

With this objective in mind, we assessed the transportability of the Tissue-Aware model trained on TCGA using only TFs across three diverse datasets: GTEx [26], TARGET [27], and the CCLE [28]. The learning procedure mirrored our previous approach: clustering TCGA samples into “Pseudo- Tissues” and training a Tissue-Aware model with TFs as input predictors. Following training, we leveraged this model to predict the transcriptomes of samples in the three aforementioned datasets using their respective TFs as predictors.

## 5. Conclusions

In this study, we demonstrated that using a small subset of transcription factors and microRNAs as selected features could effectively represent cell states and predict gene expression across 31 cancer types. Our results support the hypothesis that these regulatory molecules, due to their causal roles in gene regulation, can provide robust predictors of the transcriptome. The Tissue-Aware model, which incorporates tissue-specific information, outperformed the Tissue-Agnostic model, suggesting the importance of context for gene regulation. While our selected features were nearly as predictive as the L1000 genes, the model’s transportability was affected by the covariate shift, particularly in microRNA expression. By using only TFs in the model, we improved performance on the CCLE dataset, emphasizing the need for robust feature selection across different datasets. Future studies should explore additional regulatory molecules, such as epigenetic modifiers and chromatin remodeling complexes, as well as methods from the domain adaptation and multi-output prediction literature, to enhance our understanding of gene regulation and improve model transportability.

## Author Contributions

Conceptualization, K.R.C.; methodology, A.A. and K.R.C.; software, A.A. and K.R.C.; validation, A.A. and Z.B.A.; formal analysis, A.A.; data curation, A.A. and K.R.C.; writing—original draft preparation, A.A.; writing—review and editing, A.A., Z.B.A., H.H.P., K.R.C.; visualization, A.A.; supervision, K.R.C.; project administration, A.A.; funding acquisition, A.A. All authors have read and agreed to the published version of the manuscript.

## Institutional Review Board Statement

Not applicable

## Informed Consent Statement

Not applicable

## Data Availability Statement

The original data presented in the study are openly available in UCSC Xena at https://xena.ucsc.edu/download-data/.

---

**Figure S1.**
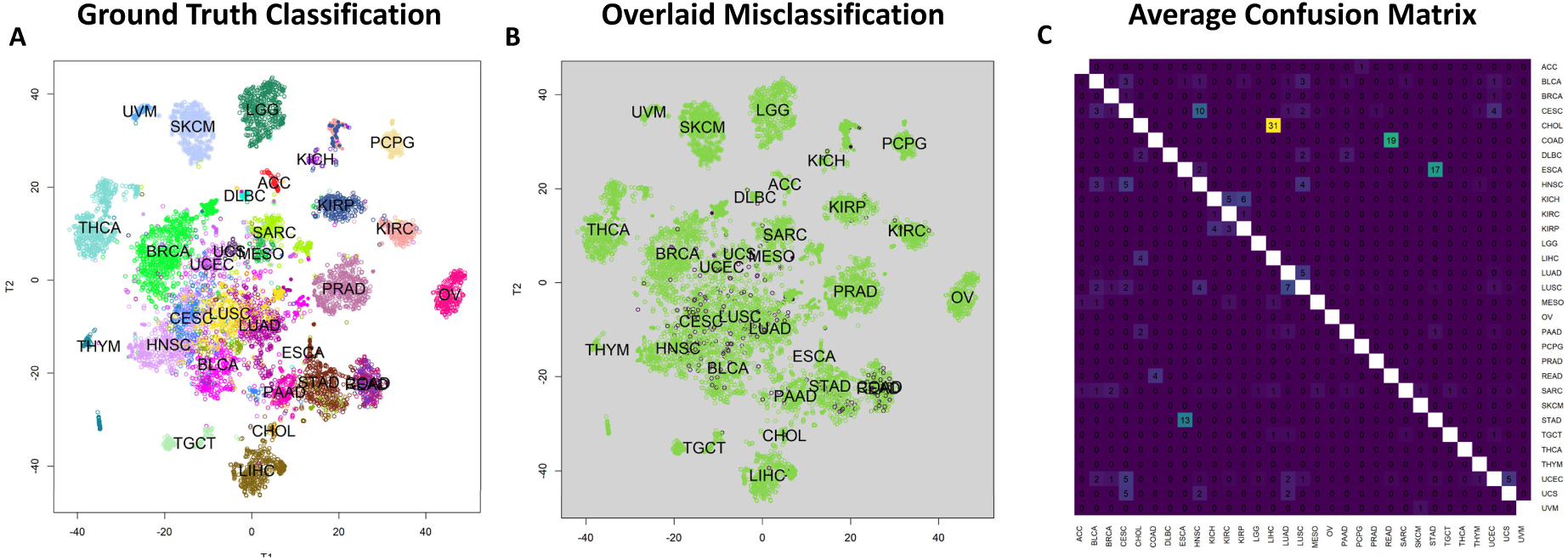
Cancer Type Classification and Visualization Through Regulatory Molecules for TPM normalized TCGA Data. **(A)** A 2D t-SNE plot of the 8895x47 input TF and miRNA expression matrix, consisting of selected 19 TFs and 28 miRNAs, with colors corresponding to known cancer types, showcasing the ability to differentiate most cancers using 47 regulatory molecules. Note that COAD (Colon Adenocarcinoma) and READ (Rectal Adenocarcinoma) appear almost superimposed due to their high similarity, causing their labels to overlap. **(B)** An enlarged t-SNE plot focusing on misclassified samples, marked in dark purple, indicates that classification errors are more common in highly similar cancer types. **(C)** An average confusion matrix generated from 10-fold cross-validation, exposing the error pattern of the SVM classifier with 90.24% accuracy. The numbers within each cell of the heatmap represent the average percentage of times, across 10-fold cross-validation, that samples from one class (row) were predicted to belong to another class (column). A value of 100% would indicate that all samples of a given class were consistently misclassified as another specific class across all cross-validation folds. Diagonal cells representing correct classifications have been left blank for clarity. Lighter matrix entries (excluding diagonals) represent larger errors, and the error pattern predominantly coincides with the t-SNE visualization of errors in panel B. Classifying similar cancer pairs like STAD and ESCA, COAD and READ, LUSC and LUAD, higher-order similar cancers such as KIRP, KIRC, and KICH, as well as squamous cancers (or cancers with a squamous cell subtype) including LUSC, CESC, HNSC, and BLCA, is challenging. All results are similar to FPKM normalized data with 56 features.

**Figure S2.**
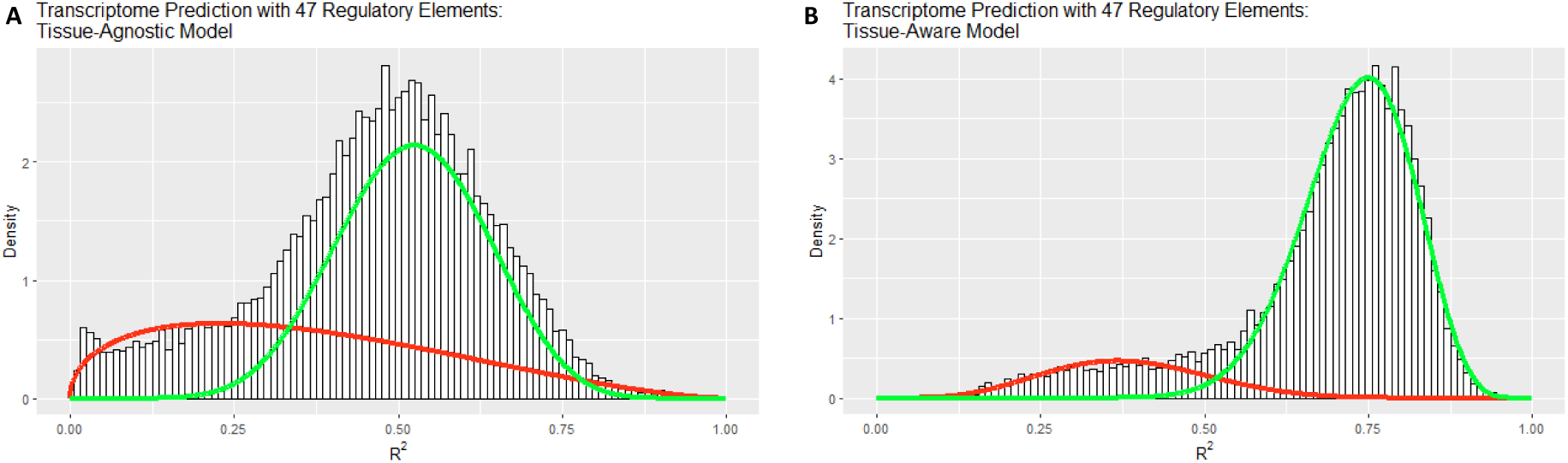
Gene Expression Prediction of Selected Regulatory Molecules with TPM Normalized Data. **(A)** and **(B)** *R*^2^ histograms for predicting 20,289 gene expressions using 8,895 samples in TCGA with Tissue-Agnostic and Tissue-Aware models, respectively, illustrating the mixture of two components: well-explained genes (green curve) and poorly-explained genes (red curve). The Tissue-Aware model demonstrates enhanced performance, with an average *R*^2^ value of 0.68 compared to 0.46 for the Tissue-Agnostic model.

**Figure S3.**
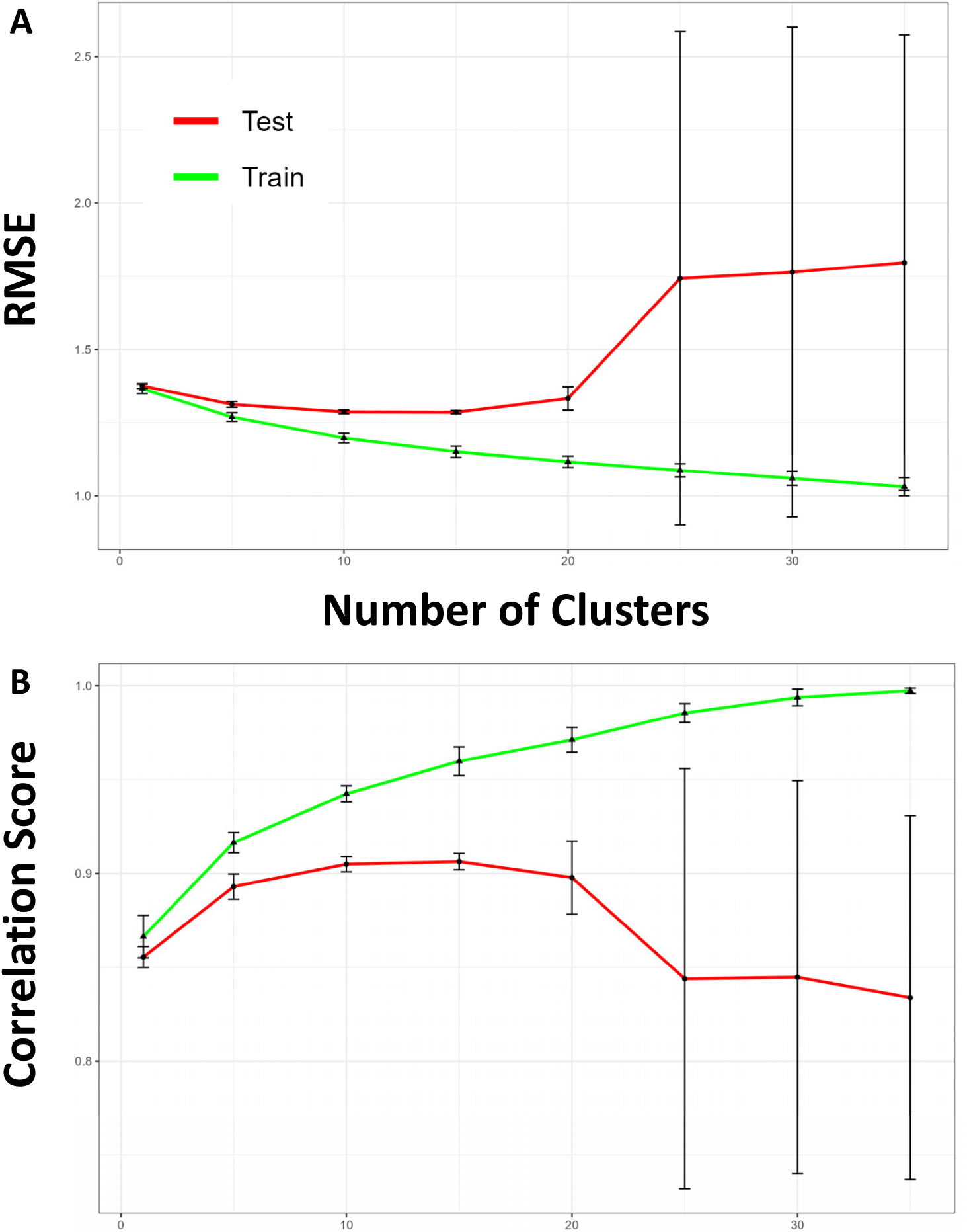
Optimizing Pseudo-Tissues for Tissue-Aware Model Generalization using TCGA TPM Normalized Data. **(A)** and **(B)** Cross-validation curves for tuning the number of Pseudo-Tissues (sample clusters) based on RMSE and correlation score, respectively, using the TCGA dataset. The optimal number of Pseudo-Tissues is determined to be 15 for both criteria. However, cross-validation assumes that the train and test data share the same distributions, which is not the case when distribution shifts are present, such as when selected miRNA expressions are heavily shifted in CCLE, thus hindering the transportability of the predictive model.

